# Single-cell RNA-seq reveals CD16^-^ monocytes as key regulators of human monocyte transcriptional response to *Toxoplasma*

**DOI:** 10.1101/863274

**Authors:** Anirudh Patir, Anton Gossner, Prakash Ramachandran, Joana Alves, Tom C. Freeman, Neil C. Henderson, Michael Watson, Musa A. Hassan

**Affiliations:** Division of Genetics and Genomics, The Roslin Institute, University of Edinburgh, Edinburgh, EH25 9RG, UK; Division of Infection and Immunity, The Roslin Institute, University of Edinburgh, Edinburgh, EH25 9RG, UK; Centre for Inflammation Research, The Queen’s Medical Research Institute, The University of Edinburgh, Edinburgh, EH16 4TJ, UK; Centre for Tropical Livestock Genetics and Health, The University of Edinburgh, Edinburgh, EH25 9RG, UK

**Keywords:** Single-cell, RNA-seq, human monocytes, *Toxoplasma*

## Abstract

Monocytes are among the major myeloid cells that respond to *Toxoplasma*, a ubiquitous foodborne that infects ≥1 billion people worldwide, in human peripheral blood. As such, a molecular understanding of human monocyte-*Toxoplasma* interactions can expedite the development of novel human toxoplasmosis control strategies. Current molecular studies on monocyte-*Toxoplasma* interactions are based on average cell or parasite responses across bulk cell populations. Although informative, population-level averages of monocyte responses to *Toxoplasma* have sometimes produced contradictory results, such as whether CCL2 or IL12 define effective monocyte response to the parasite. Here, we used single-cell dual RNA sequencing (scDual-Seq) to comprehensively define, for the first time, the monocyte and parasite transcriptional responses that underpin human monocyte-*Toxoplasma* encounters at the single cell level. We report extreme transcriptional variability between individual monocytes. Furthermore, we report that *Toxoplasma*-exposed and unexposed monocytes are transcriptionally distinguished by a reactive subset of CD14^++^CD16^-^ monocytes. Functional cytokine assays on sorted monocyte populations show that the infection-distinguishing monocytes secrete high levels of chemokines, such as CCL2 and CXCL5. These findings uncover the *Toxoplasma*-induced monocyte transcriptional heterogeneity and shed new light on the cell populations that largely define cytokine and chemokine secretion in human monocytes exposed to *Toxoplasma*.

## INTRODUCTION

Majority of lethal human pathogens spend a significant portion of their life-cycle inside immune cells, mostly monocytes and macrophages^1,2^. The intracellular lifestyle potentially enable these pathogens to not only evade the host’s immune surveillance but also antimicrobial therapy^2^. As such, a molecular understanding of pathogen encounters with host immune cells has the potential to identify novel therapeutic targets. Current knowledge on host-pathogen interactions is largely based on experiments performed in bulk populations of host and/or pathogen cells. However, host-pathogen interaction is mostly a single cell problem involving dynamic host and pathogen gene regulatory programs that often produce distinct outcomes in individual cells in a host. For example, bacteria encounters with macrophages from the same individual can simultaneously produce macrophages that; 1) are infected, while others remain uninfected, 2) kill the ingested bacteria, while others support intracellular bacterial survival, and 3) undergo programmed cell death to release the bacteria, while others survive and allow bacterial growth or persistence^3^. Similar intraindividual heterogenous infection outcomes have been observed *in vitro* during viral infections^4,5^ and *in vivo* during active tuberculosis, during which sterilized and active lesions occur simultaneously in the same host^6^. Phenotypically distinct variants of the same pathogen, such as dormant and actively replicating *Mycobacterium*, have also been isolated from the same infected host^7^. Although the molecular mechanisms underpinning these distinct intraindividual infection outcomes are largely unknown, it is plausible that they impact disease pathogenesis and antimicrobial therapy^8^. Thus, to understand the biology of infectious diseases and develop effective control strategies, it is important to consider the overall outcome of an infection as a manifestation of multiple distinct infection outcomes occurring simultaneous in individual cells within a host.

*Toxoplasma gondii*, the etiological agent for toxoplasmosis, is a zoonotic protozoan that infects virtually all warm-blooded vertebrates^9^. In human peripheral blood, monocytes are among the major myeloid cells that respond to the parasite by secreting interleukin 12 (IL12)^10^, which is required to induce the production of the indispensable anti*-Toxoplasma* interferon-gamma (IFNγ) cytokine^11^. Although usually studied in bulk host and/or parasite cells, when *Toxoplasma* interacts with monocytes from the same individual, several possible outcomes can occur to produce distinct monocyte and parasite subpopulations. The parasite can either enter the cell via active invasion to reside within a non-fusogenic parasitophorous vacuole (PV) or be taken up via phagocytosis to reside in a phagosome^10,12^. In certain cases, phagocytosed parasites can subsequently escape the phagosome to establish a PV^13^. Although not infected, some monocytes can also be exposed to, and manipulated by, secreted parasite factors through contact-dependent injection of parasite molecules^14^. Some infected monocytes can undergo programmed cell death^15^ thereby killing the parasite within, while some uninfected bystander monocytes can be activated to produce high levels of IL12^16^. There are multiple ways to achieve each of these infection outcomes, further expanding the number of molecular pathways that may be regulating the outcome of monocyte-*Toxoplasma* encounters. Additionally, among canonical human monocyte subsets (classical, CD14^++^CD16^-^; intermediate, CD14^++^CD16^+^ and; non-classical, CD14^+^CD16^+^) effective host response to the parasite, defined by the production of IL12, is reportedly restricted to CD16^+^ monocytes that phagocytose the parasite^10^. Although informative, probably due to averaging of diverse sets of cell responses, cell population-level studies on monocyte-*Toxoplasma* encounters have sometimes presented contradictory results. For example, studies on bulk human monocytes and parasite populations have presented contrasting results regarding monocyte secretion of IL12 in response to *Toxoplasma*^10,17^. To resolve such discrepancies and characterize heterogeneous monocyte-*Toxoplasma* interactions, we need to go beyond population averages and define single cell responses that when combined represent the entire monocyte and parasite responses.

Single cell RNA sequencing (scRNA-seq), which captures transcript abundance in single cells^18^, can provide the high-resolution needed to resolve transcriptional programmes underlying disparate host-pathogen interactions. scRNA-seq has so far been used to characterise, among others, the transcriptional profiles underpinning: intra-individual disparate macrophage response to *Salmonella*^3^; latent and reactivating HIV-infected human CD4+ T cells^19^ from the same donor; and the sexual commitment and development of individual *Plasmodium* parasites^20,21^. While host-pathogen interactions involve two organisms with distinct transcriptomes, scRNA-seq profiling of host-pathogen interaction is typically restricted to a single organism at a time. Single-cell dual RNA-sequencing (scDual-Seq), a hybrid of scRNA-Seq and dual RNA-seq^22^, can be used to simultaneously monitor the host and pathogen transcriptomes during an infection^23,24^.

Here, we exploit the fact that both *Toxoplasma* and human mRNA are polyadenylated and can be simultaneously profiled using standard scRNA-seq protocols to investigate the transcriptional hallmarks of *Toxoplasma* interactions with monocytes from the same donor. We report significant heterogeneity among individual monocytes and parasites. Furthermore, we find that *Toxoplasma*-exposed and control monocytes are transcriptionally distinguished by non-classical monocytes and a novel subset of reactive classical monocytes.

## RESULTS

### Defining the single-cell transcriptome of *Toxoplasma*-exposed human monocytes

In this study, we used single-cell RNA sequencing (scRNA-seq; 10X Genomics) to perform unbiased transcriptional analysis of human monocytes exposed to *Toxoplasma* for 1 hour. We conducted several pre-processing steps (Materials and Methods) including quality control, normalization, and scaling^25^ to remove potential technical bias. The processed control (unexposed) monocyte expression matrix contained 3,136 cells and 12,023 genes while the *Toxoplasma*-exposed monocyte data contained 1,352 cells and 10,795 genes. Unsupervised graph-based clustering partitioned the unexposed monocytes into two clusters; hU1 (human-unexposed) and hU2, which correspond to the non-classical (CD14^+^CD16^++^) and classical (CD14^++^CD16^-^) monocyte subsets, respectively^26^, (**Figure 1A**). The *Toxoplasma*-exposed monocytes clustered into three distinct groups: hE1 (human-exposed), which is composed entirely of non-classical monocytes; hE2; and hE3, both of which were made mostly up of classical monocytes (**Figure 1B**). The proportions of classical and non-classical cells identified in both conditions (**Figure 1A-B**) is largely consistent with reports on the composition of circulating human monocytes^27^.

**Figure 1:**
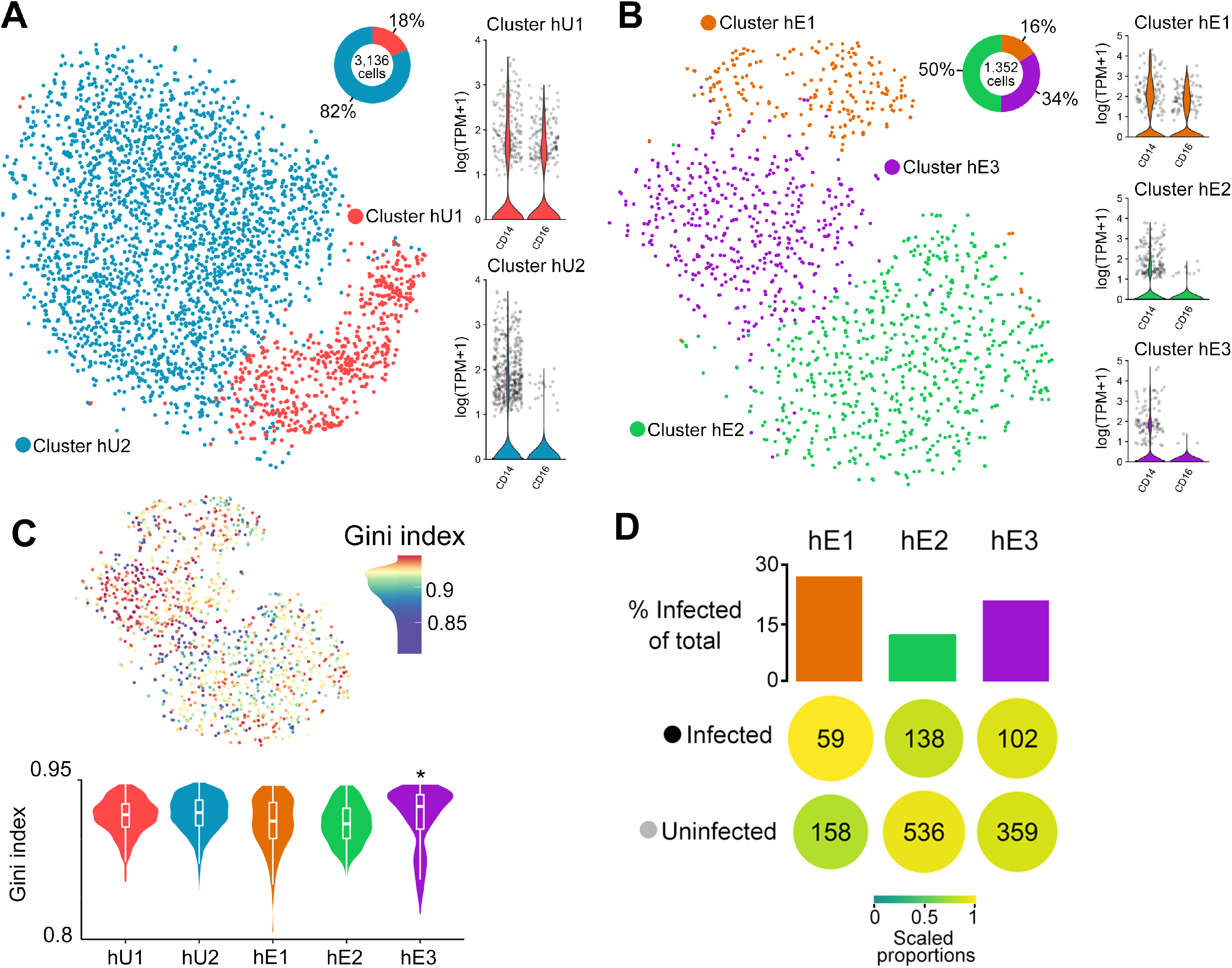
Transcriptional heterogeneity within *Toxoplasma*-exposed and unexposed human monocytes: *t*SNE plots of monocyte clusters, including non-classical and classical monocytes, of **A)** unexposed (hU1 & hU2) and **B)** *Toxoplasma*-exposed (hE1, hE2, and hE3) to human monocytes. For each condition the proportion of cells within a cluster and the expression of monocyte subset marker genes (*CD14* and *CD16*) are shown. **C)** A *t*SNE plot (Top panel) of *Toxoplasma*-exposed monocytes showing the Gini coefficients for each cell. Higher Gini index represent higher variation of expression across genes. Violin plots of the Gini coefficients for each monocyte cluster from *Toxoplasma*-exposed and unexposed monocytes (Bottom panel). **D)** The proportion of infected and uninfected cells within each cluster. Additionally, highlighting preferential infectivity of clusters (colour and size) while scaling for the total infected/uninfected cells and the number of cells within a cluster

Next we used a Gini coefficient metric, a measure of population inequality ranging from zero (complete equality) to one (complete inequality) to evaluate gene expression variability within the monocyte subsets. Cells from the hE3 cluster showed significantly (*FDR* < 0.05, medium Gini index = 0.93) higher Gini index relative to the other clusters (**Figure 1C**). In scRNA-seq mRNAs from the parasite and the monocyte it infects are tagged with the same cell barcode, which enables the reliable identification of *Toxoplasma*-exposed monocytes that are infected (contain parasite mRNA) and uninfected (do not contain parasite RNA). Thus, to determine whether the differences between monocyte clusters was due to differences in the number of infected monocytes, we examined the proportion of infected cells in each of the *Toxoplasma*-exposed monocyte clusters. There were no significant differences in the proportion of infected cells in each cluster (**Figure 1D**). However, it is worth noting that by injecting its effector proteins into host cells, *Toxoplasma* can modulate the transcriptome of host cells without truly infecting them. As such, part of what we consider exposed-uninfected (bystander) cells in this study may include cells that are injected with parasite effector proteins or infected cells from which we were not able to recover parasite mRNA. Nevertheless, when we restricted the unsupervised hierarchical clustering to the cells that contain, or do not contain parasite mRNA, we did not observe new clustering patterns (not shown). Taken together, we find that individual monocytes from the same donor respond variably to *Toxoplasma* at the transcriptional level and that this transcriptional heterogeneity is more pronounced in the classical monocyte (CD14+CD16-) subset.

### CD16^-^ monocytes transcriptionally distinguish Toxoplasma-exposed and unexposed human monocytes

Previous immunological and parasitological assays in human monocytes have reported differential response to *Toxoplasma* between classical and non-classical monocyte subsets, as well as between monocytes that are actively invaded and those that take up the parasites through phagocytosis^10,28^. Having observed transcriptional heterogeneity in the *Toxoplasma*-exposed monocytes, we determined whether a distinction between responsive and unresponsive monocyte subsets can be discerned at the transcriptional level. A *t*-distributed stochastic neighbour embedding (*t*SNE) plot, which depicts the similarity between cells based on their gene expression, of the combined *Toxoplasma*-exposed and unexposed monocytes scRNA-seq data showed that the exposed and control cells are clearly transcriptionally distinguished by the hE3 cluster (**Figure 2**). As indicated above, although hE3 distinguished *Toxoplasma*-exposed and unexposed monocytes, there was no evidence that cells in this cluster contained more parasite genes than the hE1 or hE2 clusters.

**Figure 2:**
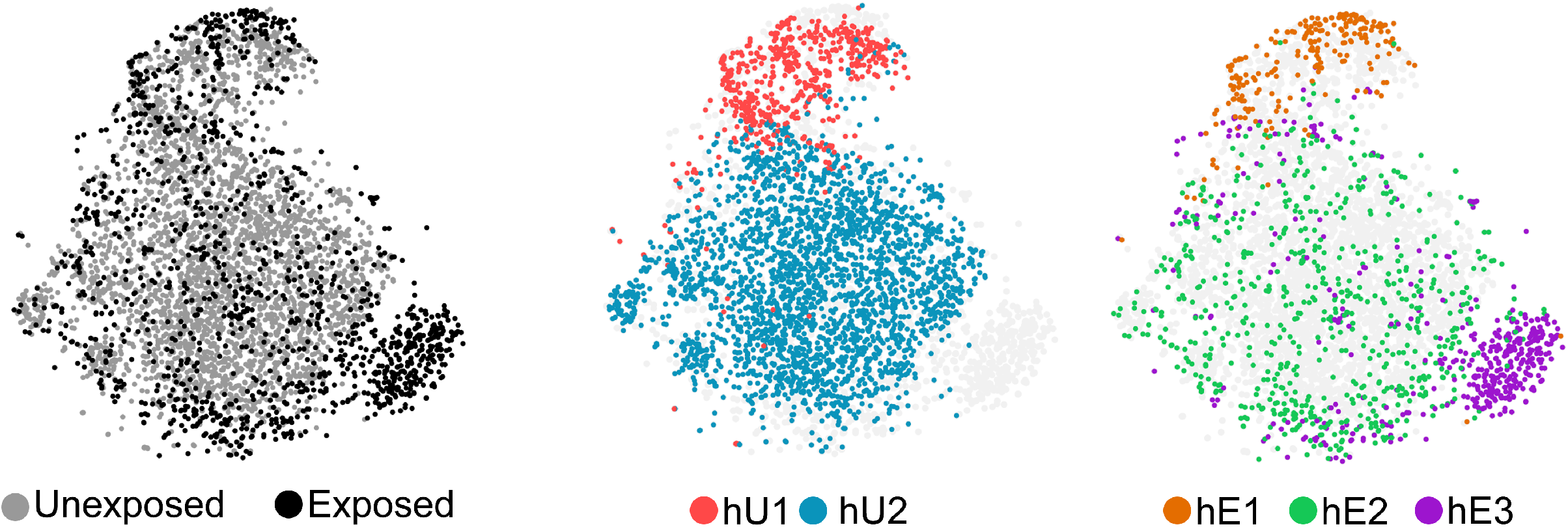
A unique subset of CD16^−^ monocytes distinguish exposure to *Toxoplasma*: **A)** *t*SNE plots of the combined *Toxoplasma*-exposed and unexposed monocytes datasets overlaid with the monocyte clusters identified from each condition (Centre and left panels).

To gain further insights into the *Toxoplasma*-exposed monocytes, we sub-divided cells in each cluster into infected (contain parasite mRNA) and uninfected (no parasite mRNA) and preformed differential expression (DE) analysis between the clusters. 711 genes were differentially expressed (**S1A**), of which 418 were highly expressed in specific monocyte clusters (**Figure 3A**). 218 genes were highly expressed in the hE3 cluster, of which 22 were highly expressed in the infected and 196 in both the infected and uninfected monocytes (**Figure 3A** and **S1A**), suggesting that the transcriptional heterogeneity in the hE3 cluster is driven by both the infected and uninfected bystander cells. Interestingly, the number of human genes and reads were significantly lower (FDR < 0.05) in hE3 infected and uninfected cells relative to other groups (**Figure 3B**), without a significant difference in the number of parasite genes. Similarly, out of the 80 highly expressed genes in the hE1 cluster, 61 (76%) were expressed in both the infected and uninfected cells. Despite being like the hE3 cluster in monocyte subset composition, the hE2 cluster had the least number (16) of highly expressed DE genes.

**Figure 3:**
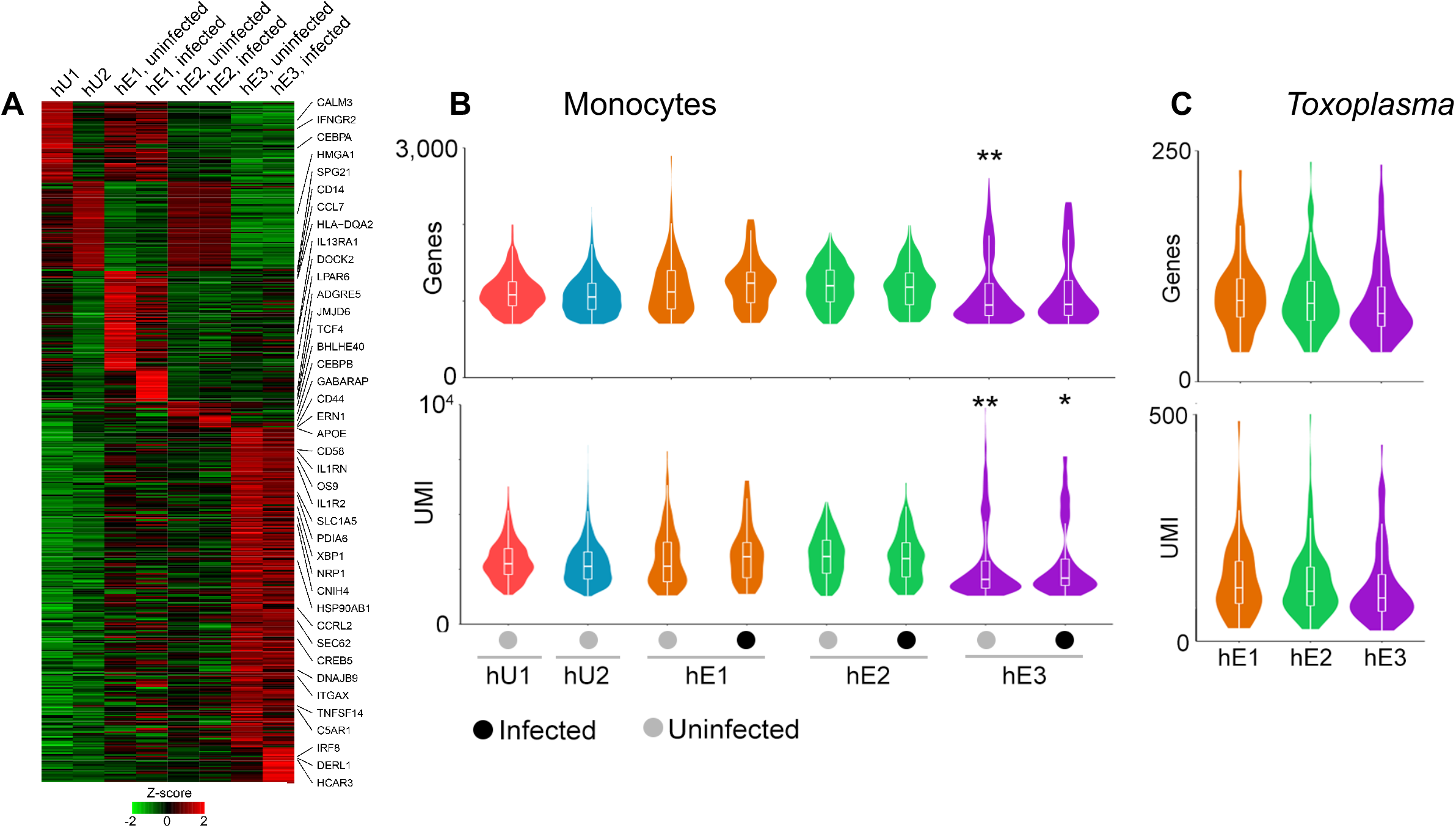
Gene expression across monocyte subsets: **A)** A heatmap showing the average expression of the 418 DE genes for each monocyte cluster in *Toxoplasma*-exposed and unexposed monocytes. The number of genes and reads mapped to **B)** Monocytes and **C)** invading *Toxoplasma*. *Significance at *FDR* < 0.05, ** significance at *FDR* < 0.01.

To functionally annotate DE genes in hE1 or hE3 clusters, enrichment analysis was performed. The top Gene Ontology terms enriched in DE genes that were highly expressed in both the uninfected and infected cells in hE3 cluster included innate immune associated terms, such as “immune effector process” (*FDR* = 3.48×10^−4^), “myeloid cell activation involved in immune response” *(FDR =* 4.13×10^−7^), and “cell activation involved in immune response” (*FDR* = 2.95×10^−6^) (S1B). Among the DE genes in the hE3 cluster were pro-inflammatory chemokines that are known to be induced in human peripheral blood mononuclear cells and monocytes exposed to *Toxoplasma*^17^, such as *CXCL8*, (for infected, fold change = 1.5, *FDR* = 2.11×10^−3^; uninfected, fold change = 2.9, *FDR* = 7.61×10^−59^). GO terms enriched in the uninfected cells in the hE3 cluster included “response to endoplasmic reticulum unfolded protein” (*FDR* = 9.91×10^−8^) and “endoplasmic reticulum part” (*FDR* = 6.45×10^−15^). Consistent with the GO enrichment analysis, the x-box binding protein (*XBP1*), a transcription factor that is critical for the induction of genes required for resolving endoplasmic reticulum stress induced by the accumulation of unfolded proteins^27^, was one of the highly expressed DE genes in the uninfected cells in hE3 cluster (**Figure 3A**). Besides *XBP1*, DE genes highly expressed in the uninfected cells in hE3 included other unfolded protein response (UPR)-related genes such as *DNAJB9* (a target of *XBP1*), *OS9*, and *PDIA6* (**Figure 3A**). GO terms enriched in the infected cells in the hE3 cluster included “cellular response to LPS” (*FDR* = *5.05E-2*) and “cellular response to molecules of bacterial origin” (*FDR* = *5.58E-2*). Enrichment analysis for the hE1 cluster also revealed immune response associated terms such as, “regulation of immune system process” (FDR = 7.53×10^−4^) and “defence response” (FDR = 7.71×10^−3^). The top cellular component terms enriched in the uninfected cells in the hE1 cluster included “lysosome” (*FDR* = 1.5×10^−4^) and “lytic vacuole” (*FDR* = 1.5×10^−4^). Congruent to observations in the THP-1 cell line (a human monocyte cell line that replicates most *Toxoplasma* infection phenotypes observed in primary human monocytes) that are infected or separated from *Toxoplasma* in transwell^17^, *CCL2* was DE in both the infected (1.9-fold, *FDR* = 7.57×10^−3^) and uninfected (3.3-fold, *FDR* = 3.35×10^−17^) cells in the hE1 cluster (S1A). The expression of *CCL2* in *Toxoplasma*-exposed monocytes is reportedly initiated by the S100 calcium-binding protein A11 (S100A11)^17^. However, unlike the other members of the S100 family, including *S100A4, 6* and *9* that were DE in hE1, *S100A11* was not. Considering all DE genes, those highly expressed in hE1 included members of the MHC class 2 family and genes associated with interferon response (IFIT, *IRF* and *OAS* gene families). Though few genes were DE solely in hE2, the cluster showed a high expression of immune related genes that were also expressed in other groups, including *CCL3, CCL4, NFKBIA* and *IL1β*. Certain DE genes expressed in hE3 and hE1 such as members of the MHC class 1, cathepsins and proteasome complex, were not expressed in the hE2 cluster. In summary we reveal an unprecedented level of transcriptional heterogeneity in *Toxoplasma*-exposed monocytes and that transcriptional response to the parasite at 1 h post exposure is defined mostly by a subset CD16^−^ monocytes.

### Monocyte response to *Toxoplasma* is defined by chemokine secretion

The observation that a subset of CD16^−^ monocyte subset (hE3) transcriptionally distinguish *Toxoplasma*-exposed and unexposed monocytes prompted us to investigate whether the transcriptional heterogeneity translate into a distinctive infection phenotype. To explore this, we used monoclonal antibodies against the cell surface protein product of the *CD98* gene that is highly expressed in the hE3 cluster to sort *Toxoplasma*-exposed monocytes followed by functional assays. We used a human multiplex cytokine array to measure the level of several cytokines and chemokines, including CCL2 and IL12, in cell-free supernatants from the sorted CD98^+^ monocytes (**Figure 4A**). CD98^−^ monocytes were used as controls. The sorted CD98^+^ cells secreted significantly more CXCL5, CCL2 and MIP1, but less IL12 and IL6, compared to CD98^−^ monocytes (**Figure 4B**). The level of IL1β, known to be secreted by monocytes in response to *Toxoplasma*^15^, was induced in CD98^+^. Therefore, *Toxoplasma*-induced transcriptional heterogeneity observed in human monocytes can be linked to specific Toxoplasma-induced monocyte phenotypes at the protein level.

**Figure 4:**
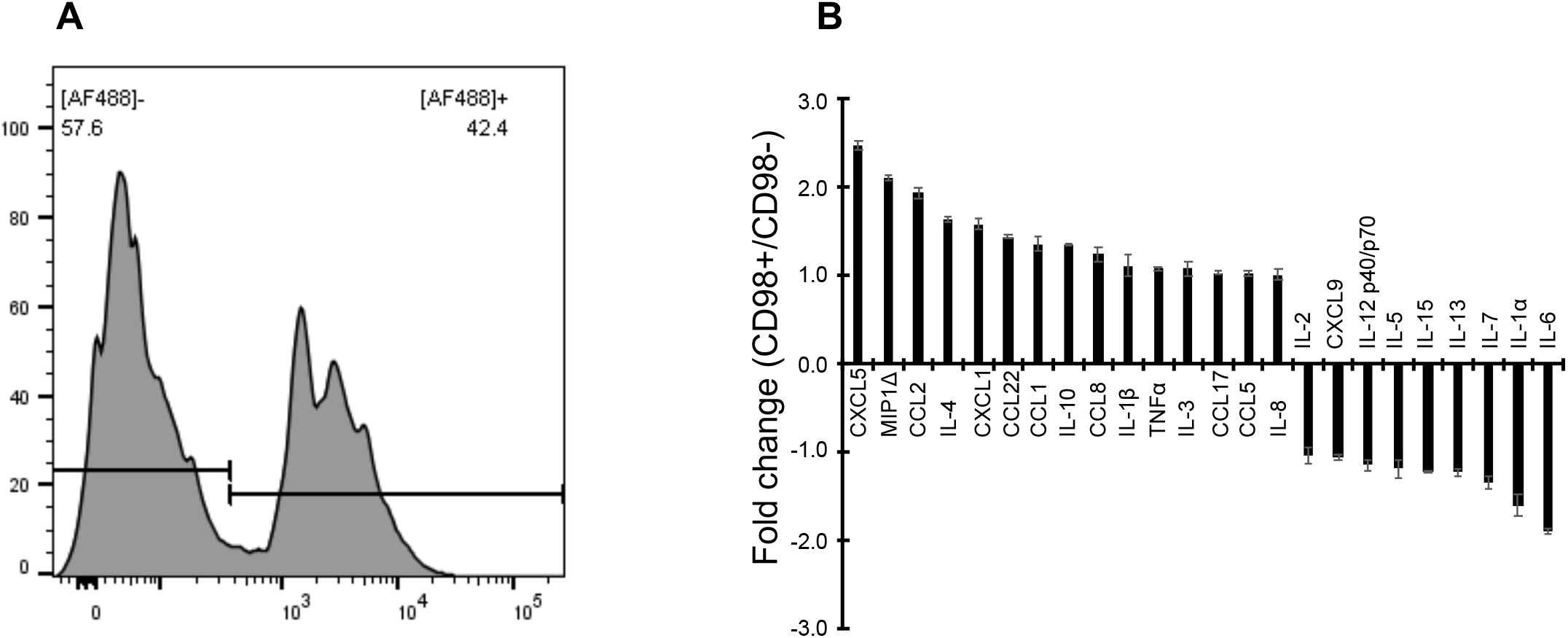
A functional analysis of infection-distinguishing *Toxoplasma*-exposed monocytes: **A)** At 1 h post-infection, the percentage of CD98^+^ (Alexa Flour 488, AF 488) monocytes were sorted for functional assays. **B)** Fold changes of cytokine and chemokine levels in the sorted CD98^+^, relative to control (CD98^−^) monocytes. Values are mean ±s.e of three independent biological repeats.

### Transcriptional heterogeneity of *Toxoplasma* exposed to monocytes

Host-pathogen encounters are highly dynamic processes modulated by both host-and pathogen-derived factors. Therefore, we investigated whether parasites display different transcriptional profiles in individual monocytes. After data pre-processing and quality control we recovered 873 parasites, of which 299 shared cell barcodes with individual monocytes. In total, the 299 *Toxoplasma* cells expressed 2,556 genes. We initially grouped parasites based on the type of monocyte they infected, however no DE genes were identified for any of these groups. Subsequently, parasites were clustered into three distinct groups (hT1, hT2 and, hT3) based on their gene expression (**Figure 5A**). Varying abundance of parasites were observed between clusters, the largest being hT1 (585 parasites) and the smallest hT3 (109 parasites) (**Figure 5B & C**). However, in all cases approximately the same proportion, 33% to 37% of parasites were linked to monocytes. To determine whether there were differences between infecting and bystander parasites for a given cluster, the number of genes and reads were examined for each. Significant differences were observed between infecting and bystander parasites of hT1 and hT2 (**Figure 5D**). However, on conducting DE analysis between the infecting and bystander parasites for each cluster no DE genes were identified. To identify the genes that underpin the parasite transcriptional heterogeneity in individual monocytes, we performed DE analysis between the parasite clusters. Of the 169 DE genes 40, 73, and 80 were highly expressed in hT1, hT2 and hT3, respectively (**S2A**). The top DE genes included several ribosomal proteins, the microneme protein 10 (*MIC10*), a putative elongation factor 1-alpha (*EF-1-ALPHA*), and a dense granule protein *(GRA12)* in hT1; SAG-related sequence (*SRS20A*), several rhoptry and rhoptry neck proteins such as *ROP17* and *RON8* in hT2 and; a microtubule-associated protein (*SPM1*), *GAP45*, and several hypothetical proteins in hT3. Next, we analysed the biological processes enriched in the DE genes using functional gene enrichment analysis based on the gene ontology terms available in ToxoDB^30^. The hT1 cluster, in which several ribosomal genes were DE, was enriched in “translation” (*FDR* = 1.6×10^−7^) and “peptide biosynthetic processes” (*FDR* = 1.6×10^−7^) (**S2B**). The hT2 and hT3 clusters were enriched in “Apical part of cell” and “Pellicle” terms associated cellular localization (*FDR* < 0.001). In summary, these results show that individual parasites transcriptionally respond to the monocytes they infect. However, from the current study, it is not apparent that the parasite transcriptionally respond to the cognate monocyte transcriptional landscape.

**Figure 5:**
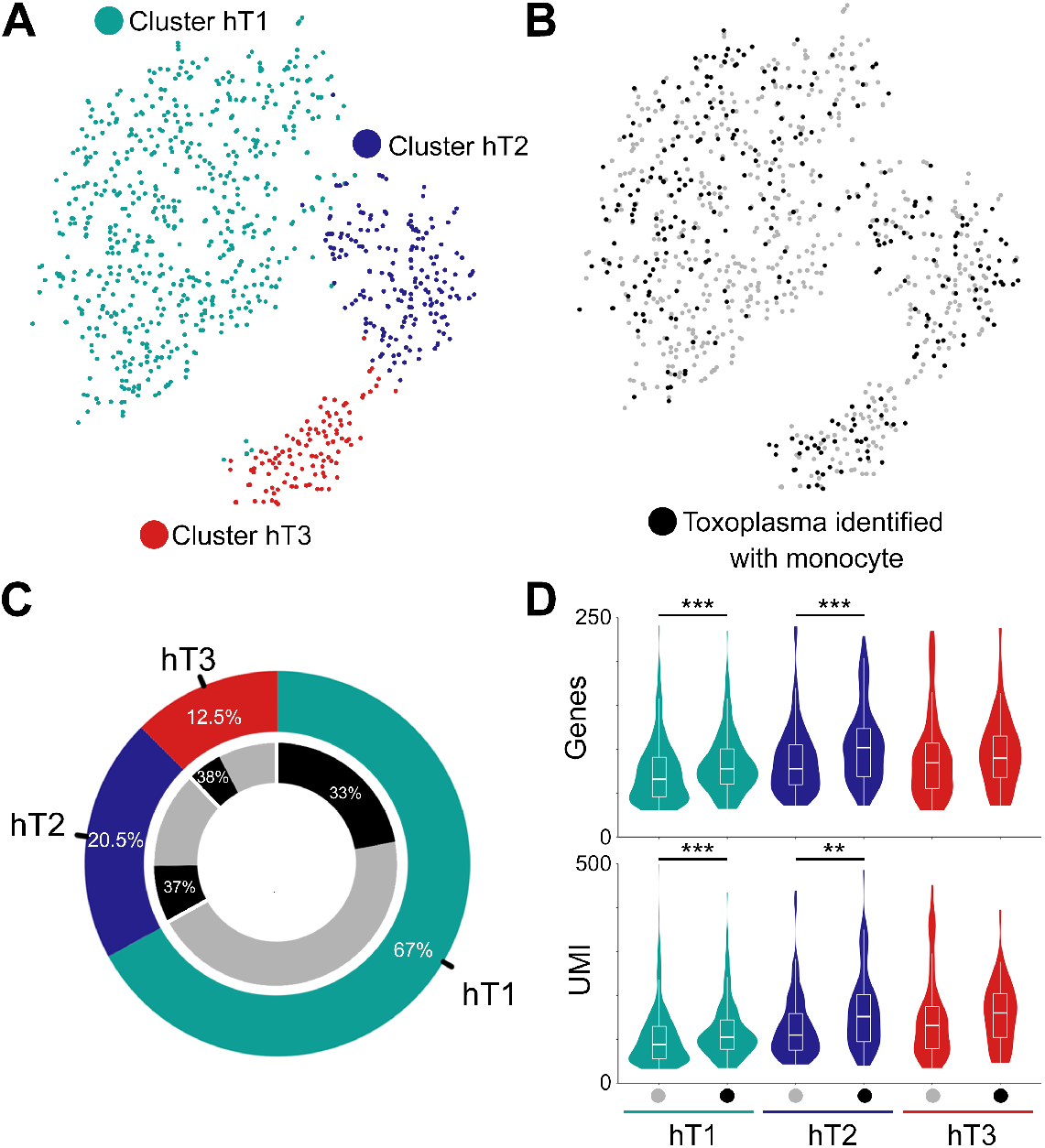
Transcriptional heterogeneity of *Toxoplasma* exposed to monocytes: **A-C)** *t*SNE and clustering of *Toxoplasma* parasites with the proportion of cells shown for each cluster (hT1, hT2 and hT3). D) The distribution of reads and *Toxoplasma* genes expressed based on their clustering and whether they infected (black dot) a monocyte. ** Significance at *FDR* < 0.01, *** significance at *FDR* < *0.001*.

## MATERIAL AND METHODS

### Blood donors and monocyte isolation

Human whole blood was collected from healthy adult donors who provided written informed consent. Blood was collected according to the guidelines of and with approval from the Roslin Institute’s Health and Safety committee. Monocytes were isolated from peripheral blood mononuclear cells as previously described^10^, to yield >98% pure monocytes.

### Parasites and infection

RH strain tachyzoites, maintained by serial passage on human foreskin fibroblasts (HFFs; originally obtained from the Boothroyd lab, Stanford University), were used in all infections. Parasites were grown in RPMI (Life Technologies) supplemented with 1% foetal bovine serum (FBS; Omega Scientific), 2 mM glutamine (Sigma), 10 mM HEPES (pH 7.5; Sigma), and 20 μg/ml gentamicin at 37°C in 5% CO_2_. The parasites used for monocyte infection were prepared by scraping T-25 flasks containing heavily vacuolated HFFs followed by sequential passage through 25G and 27G needles. The released parasites were pelleted by centrifugation at 572 × g for 7 min, washed in phosphate-buffered saline (PBS; Life Technologies), filtered using a 5 μm membrane to exclude host cell debris, and counted. 2 × 10^6^ monocytes were plated in 6-well tissue culture plates overnight prior to infection. The cells were left unexposed or exposed to freshly prepared parasites at a multiplicity of infection (MOI) of 1:1, briefly centrifuged to bring the monocytes and parasite into contact and incubated at 37°C in 5% CO2 for 1h before processing the cells for single-cell sequencing or flow cytometry.

### Single cell RNA-sequencing

*Toxoplasma*-exposed and unexposed monocytes from one donor were harvested and washed three times in cold PBS supplemented with 0.1% bovine serum albumin (BSA, Thermo Fisher). The scRNA-seq libraries were generated using the Chromium Single Cell 3’ Library & Gel Bead Kit v.2 (10x Genomics) according to the manufacturer’s protocol. Briefly, 1×10^5^ viable cells were FACS-sorted, washed once in cold PBS, counted on the Countess II Automated cell counter (Thermo Fisher), and used to generate single-cell gel-beads in emulsion. The gel-beads in emulsion were disrupted after reverse transcription and the barcoded cDNA isolated and amplified by PCR. The resulting PCR products were fragmented, followed by end-repair, A-tailing and, adding sample indexes. The single-cell libraries were sequenced at a depth of 1 million reads per cell at the University of Edinburgh Genomics core facility on an Illumina NovaSeq machine.

### Single-cell RNA-sequencing analysis

Alignment, filtering, barcode and unique molecular identifier (UMI) counting were performed using Cell Ranger v.2.1.0 (10x Genomics) based on the human (GRCh38, Gencode) and *Toxoplasma* (ToxoDB-39) genomes. The UMI count matrices from exposed and unexposed conditions were processed seperately using the Seurat package v2^31^. Briefly, monocytes expressing less than 200 genes and with 8% mitochondrial genes were excluded. Similarly, genes detected in <10 cells were excluded from further downstream analyses. Raw UMI counts were log-normalized and cells having a total normalized expression value three standard deviations from the mean were excluded. Additionally, cells outside the 95% confidence interval of the total normalized UMI vs. number of genes per cell were filtered out. Based on the remaining samples, genes were scaled to remove the effects of library size and percent mitochondrial reads. Dimensionality reduction of the filtered genes was performed using the most significant principle components (PCs) based on Jackstraw permutations. The number of significant PCs varied with monocyte conditions, with 21 and 17 PCs in the unexposed and *Toxoplasma*-exposed monocytes, respectively. Clustering was performed using the Smart local moving (SLM) algorithm^32^. To generate the *t*SNE plots of the combined exposed and unexposed monocytes data, the 18 significant PCs from the combined dataset were used, which were based on the normalized and scaled expression matrix of all cells filtered for each condition.

*Toxoplasma* cells were filtered based on a minimum expression of 30 genes, while genes expressed in less than 3 cells were removed. Subsequently this data was normalized, scaled and significant PCs calculated. The 5 most significant PCs were used in dimensionality reduction of the *Toxoplasma* gene expression data. Functional enrichment for differentially expressed genes were performed either using ToppFun from the ToppGene suite^33^ or ToxoDB ^30^.

### Cell sorting

2 × 10^6^ monocytes from three donors were separately seeded in 6-well cell culture plates and allowed to settle overnight. Freshly prepared parasites were then added to the cells at a MOI of 1, briefly centrifuged, and incubated for an additional 1 h at 37°C with 5% CO_2_. The cells were collected, washed once with PBS, resuspended in blocking buffer (PBS supplemented with 1% Goat serum) and incubated on ice for 30 min. The cells were centrifuged at 400 × g for 4 min at 4°C and washed once in ice-cold FACS buffer. CD98^+^ were sorted in BD FACSAria III (BD Biosciences) after staining with Alexa Flour 488 anti-human CD98 (Novus Biologicals) monoclonal antibodies. Live cells were sorted after staining with the Zombie Violet viability dye (Biolegend).

### Cytokine and chemokine measurements

Sorted CD98^+^ and CD98^−^ monocytes were seeded in fresh media and incubated for 4 h before collecting cell-free supernatants, which were stored at −80°C until use. Cytokines and chemokines, including IL1β, B, IL12 and CCL2, were measured in supernatants using a multiplex Human Cytokine Array C3 kit (eBioscience) according to the manufacturer’s protocol.

### Statistical analysis

Differential expression of genes across groups of monocyte and *Toxoplasma* subsets was done using Wilcoxon test. Comparison of the number of genes or reads between groups was conducted using a one-way ANNOVA, following a post-hoc test. To identify any enrichment in monocyte or *Toxoplasma* subsets, the Fischer’s exact test was used. Where there was multiple testing, adjusted FDR was used.

## DISCUSSION

Herein, we applied a combination of scRNA-seq and phenotyping of sorted cell populations to provide a high dimensional insight into human monocyte-*Toxoplasma* interactions. We have shown that individual monocytes from the same donor exhibit great transcriptional heterogeneity in response to the parasite and that *Toxoplasma*-exposed and unexposed cells can be distinguished transcriptionally by a subset of CD16^−^ monocytes, composed of both infected and potential bystander cells. In addition, our results show that the transcriptional heterogeneity translates into functional differences between individual monocyte clusters, including differences in CCL2 and IL1β secretion, which previous studies have reported to be induced in human monocytes exposed to either *Toxoplasma* or cell-free supernatant from *Toxoplasma-infected* cells^15,16^.

Previously, CD16^+^ monocytes were reported to distinguish responsive monocytes, based on IL12 secretion^9^. Here, although CD16+ cells can transcriptionally distinguish *Toxoplasma*-exposed and unexposed monocytes, a clear distinction is only achieve with the by CD16^−^ subset. This discrepancy may be due to several factors. Besides averaging responses in cell populations, which can mask cell-to-cell differences in IL12 secretion, this and the previous study are based on different phenotypes; transcript and protein abundance, which often do not match^32^. Importantly, we investigated monocytes exposed to *Toxoplasma* for 1 hour, rather than 24 h as in the previous study^9^. Thus, it is plausible that early (1 h) responses to the parasite are defined mostly by CD16^−^ monocytes while long-term (24 h) responses are modulated by CD16+ monocytes. As such discrepancy between this and the previous study^9^ could be temporal, rather than functional. Nevertheless, we observed more IL12 in cell-free supernatants from control, compared to sorted hE3 cells, suggesting that significant differences in IL12 secretion are discernible at 1 h postinfection. Noteworthy, primary human monocyte response to *Toxoplasma* was recently reported to be defined more by the secretion of chemokines, including CCL2, than IL12^16^. Consistent with this observation, we observed differential expression of several chemokines between the different *Toxoplasma*-exposed monocyte clusters. Furthermore, sorted monocyte populations that distinguish *Toxoplasma*-exposed and unexposed monocytes secreted more chemokines, including CCL2 than the corresponding controls. We propose a future temporal scRNA-seq analysis of monocyte-*Toxoplasma* encounters to determine whether additional monocyte clusters that distinguish *Toxoplasma*-exposed and unexposed monocyte clusters emerge over time.

*Toxoplasma* transcriptional heterogeneity is consistent with observations in other intracellular pathogens infecting cells of the monocyte/macrophage lineage, including *Salmonella*^23^. *Toxoplasma* can infect phagocytic cells either via active parasite invasion or phagocytic parasite uptake, with the parasite ending up initially in the PV or phagosome^10,12,13^. Monocyte responses to the parasite, including IL12 secretion, is reportedly influenced by the route of parasite entry; only phagocytosed parasites induce IL12 secretion^10^. Although we observed transcriptional segregation within parasite-containing monocyte population, we did not observe an overrepresentation of parasite-containing cells in individual monocyte clusters (**Figure S1**). This suggests that active invasion or phagocytic parasite uptake probably does not define the monocyte transcriptional segregation or that *Toxoplasma* transcriptional heterogeneity is not modulated by intracellular niche (PV or phagosome). Currently, we lack insight into what role the route of parasite entry play in the monocyte transcriptional segregation, mostly due to a lack of well-defined transcriptional markers of actively invaded and monocytes that phagocytose the parasite. However, we are in the process of generating transcriptional profiles of sorted bulk populations of actively invaded, phagocytic, and truly bystander monocyte subpopulations, which will greatly improve our current analysis. Majority of genes that were differentially expressed in *Toxoplasma*-exposed relative to unexposed monocytes were from uninfected cells. This may suggest that the responses in bystander cells drive the phenotypic outcome of infection rather than transcriptional responses in the parasites and monocytes they infect. However, as indicated above, since we identify infected cells by the presence of parasite mRNA in it, it is possible that some of the exposed-uninfected (considered bystander) monocytes are indeed infected, but we were not able to detect parasite mRNA due to technical limitations or are truly uninfected but are injected with *Toxoplasma* effector proteins. A more in-depth analysis will be possible once we transcriptionally define different subpopulations of *Toxoplasma*-exposed monocytes, including actively invaded, phagocytic, and exposed-uninfected subpopulations.

## ACKNOWLEDGEMENTS

The authors thank the Baillie lab at the Roslin Institute for help with the blood sampling approvals and protocols. We also wish to thank the University of Edinburgh Genomics Core for providing facilities and services. MAH is funded by a University of Edinburgh Chancellor’s Fellowship and a Bill and Melinda Gates Foundation award to the Centre for Tropical Livestock Genetics and Health (OPP1127286). The Roslin Institute receives strategic investment funding from the Biotechnology and Biological Sciences Research Council. JPS was funded by a National Institute of Health grant (NIH R01AI080621).

## TABLE LEGENDS

**S1: Differential expression and enrichment analysis for monocyte subsets: A)** Differentially expressed genes in monocytes groups (infected or uninfected) within each cluster. **B**) GO terms (biological processes & cellular localization) enriched in genes DE expressed in infected or uninfected cells within each monocyte cluster

**S2 Differentially expression and enrichment analysis for *Toxoplasma* clusters: A)** Differentially expressed genes in individual *Toxoplasma* clusters. **B**) GO terms enriched in the DEGs.

**Figure S1:**
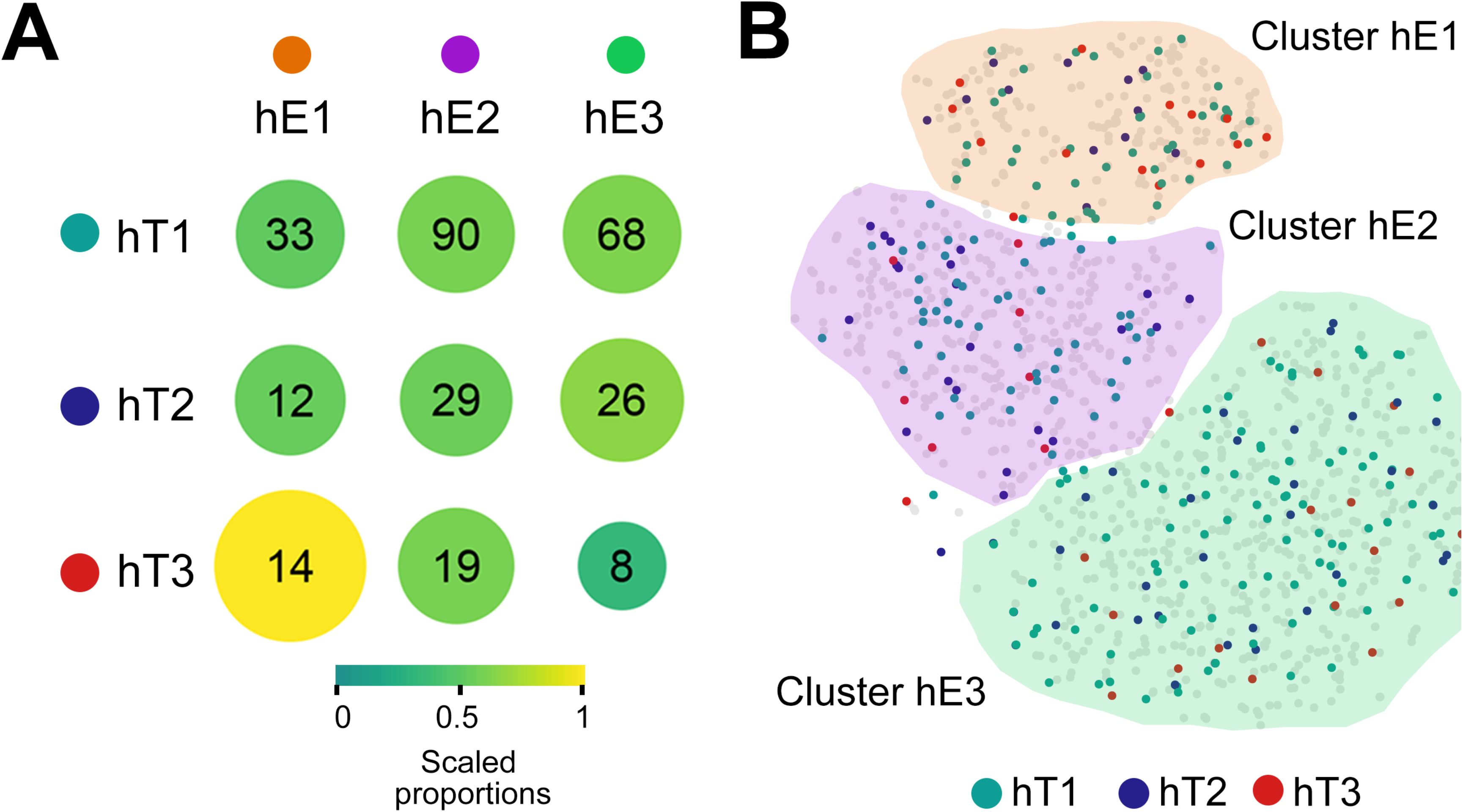
Distribution of parasites in each *Toxoplasma* cluster across monocyte subsets: **A**) The distribution of *Toxoplasma* clusters across monocyte subsets, also shown in **B**) *t*SNE plot of *Toxoplasma*-exposed monocytes.

